# A Method for Checking Recombinant Protein Quality to Troubleshoot for Discordant Immunoassays

**DOI:** 10.1101/2021.06.02.446753

**Authors:** Thi K. Tran-Nguyen

## Abstract

Discordant results of recombinant protein-based enzyme-linked immunosorbent assays (ELISA) and other antigen detection tests are a common and vexing problem in scientific research and clinical laboratories. The reproducibility of immunoassays based on antibody specificity can be adversely affected by cryptic changes in the composition of the analyte (e.g., the protein antigen). By use of sodium dodecyl sulfate-polyacrylamide gel electrophoresis and mass spectrometry analyses, we found varying levels of purity and the extent of post-translational modifications (PTMs), particularly of N-linked glycosylation and phosphorylation, among varied lots of recombinant glucose-regulated protein 78 (GRP78) expressed in *Escherichia coli.* We expect these lotto-lot variabilities of both purity and PTMs led to inconsistent results stemming from ELISA assays that measured GRP78 autoantibody levels in patient plasma specimens. We present these analyses to draw readers’ attention to a potentially common, yet seldom appreciated problem, in laboratory assays using recombinant proteins as antigens for antibody detection, and propose a workflow to detect and troubleshoot for this typical problem.

## 1. Introduction

Poor reproducibility for recombinant protein-based assays can create many challenges for both basic and clinical research, and these challenges deserve more attention to ensure the quality of both endpoint and downstream studies [1–3]. Although recombinant proteins are essential tools in laboratory assays, quality control (QC) of these reagents is often overlooked [4]. Without the proper QCs, results derived from these assays may otherwise lead to spurious and/or irreproducible data that waste time and effort, and could potentially lead to therapeutic errors [4].

Recombinant protein-based enzyme-linked immunosorbent assays (ELISA) are widely used to detect antigen-specific antibodies in serum/plasma, and other specimens, for diagnosis and monitoring of autoimmune diseases [5,6]. We have developed an indirect ELISA using recombinant Glucose-Regulated Protein 78 (rGRP78), a member of the heat shock protein family, to quantify and examine the role of GRP78 autoantibodies in chronic obstructive pulmonary disease (COPD) [7–9]. We noticed discordant results of anti-GRP78 IgG ELISA that used different lots of commercial rGRP78. We then decided to troubleshoot these varying ELISA results by examining the Coomassie Colloidal stained rGRP78 antigen in sodium dodecyl sulfate-polyacrylamide electrophoresis (SDS-PAGE) gels, and conducting nano-liquid chromatography mass spectrometry (nLCMS2) analyses to ascertain purity and identify post-translational modifications (PTM’s) and degradation products associated with the various acquired rGRP78 lots.

This approach is often referred to as a gel-enhanced liquid chromatography tandem mass spectrometry (GeLC-MS2) application, which can include either 1-dimensional (1-D) or 2-D SDS-PAGE stained gels, with or without the use of tandem mass spectrometry (MS2) for quality control (QC) purposes. The straightforward procedure has been applied in the general antibody QC workflow for a number of years [10–13], which includes the QC of specific PTM’s, predominantly glycosylation of antibody biologics [14–16].

In this project, we have undertaken a simplified and affordable approach that could be similarly adopted by any lab that has access to a MS core facility. As an example, we took commercially available 1-D gels combined with standard MS2 analysis that included proteomic searches, such as cleaved N-glycation sites following Peptide-N-glycodidase F (PNGase F) treatment (i.e. deamination products on glutamine and arginine); mass shifts associated with citrullination of arginine, unexpected cleavage sites, common oxidation events, and O-Linked ß-N-acetylglucosamine (O-GlcNac); and phosphorylation changes on serine, threonine, and tyrosine [17,18]. Utilizing the clear overview illustrated herein, one can run a gel for the first QC step, and submit a gel piece for MS2 analysis with any MS core facility.

*Escherichia coli (E. coli)* is the most commonly utilized expression system for the production of recombinant proteins. Although protein PTM is a fundamental and ubiquitous process among all living organisms [19–21], it was widely believed that *E. coli* does not possess PTM capacity. However, recent studies have reported the existence of analogous processes in prokaryotes, particularly in *E. coli* [22–24]. Advances in MS-based proteomics and analysis software have enabled us to identify the presence and quantify the abundance of a particular PTM with high confidence [25]. In this study, besides identifying the varying levels of rGRP78 purity, we also discovered different levels of PTM’s in differing lots of this commercially supplied product.

PTMs potentially cause many unwanted effects in recombinant proteins [26–28] such as improper folding, aggregation, and immunogenicity, which can lead to poor efficacy and toxicity of biosimilar drugs [26–29]. However, we found fewer reports describing similar problematic effects of PTMs in recombinant protein products used in pre-clinical or basic research immunoassays. To the best of our knowledge, the varying level of purity and PTMs of recombinant protein from commercial vendors as potential sources of inconsistent results in laboratory assays such as ELISA has not been well-documented, and many researchers may be unaware of this issue. In this report we illustrate a simplified experimental GeLC-MS2 workflow, including an informatics approach as an additional tool to QC recombinant proteins, in order to sidestep and/or troubleshoot downstream problems. We hope to raise awareness of this potential complication of recombinant protein-based assays, while helping both clinical and basic scientists along the way.

## 2. Materials and Methods

### 2.1. Commercial Recombinant GRP78 (rGRP78)

Recombinant Human GRP78 protein (Catalog No. SPR-119) were purchased from StressMarq Biosciences (Victoria, BC, Canada). Four different lots (lot#1101, lot#120914B, lot#130312 and lot#140923) of the same product obtained from this vendor from September 2014 -to-October 2016 were evaluated by SDS-PAGE and MS analyses.

### 2.2. rGRP78 Protein Expression in E. coli

Full-length human GRP78 (https://www.uniprot.org/uniprot/P11021) was cloned into the pDB-HisGST vector (DNASU, Murrieta, CA) which contains an N-terminal hexahistidine tag for downstream affinity purification. *E. coli* strain BL21(DE3) (Thermo Fisher, Waltham, MA), cultured in Luria-Bertani medium, was used as the recombinant host to express human GRP78. Following bacterial cell lysis, centrifugation, and washing, the rGRP78 was purified using nickel columns [30].

### 2.3. COPD and Smoke Control Plasma Specimens

Chronic Obstructive Pulmonary Disease (COPD) and Smoke Control (SC) subjects were enrolled from the Pittsburgh COPD Specialized Center for Clinically Oriented Research (SCCOR) cohort [31]. Inclusion criteria was a minimum 10 pack-year smoking and exclusion criteria were diagnoses with other lung and/or cardiovascular diseases in the past year, prior thoracic surgery, and a body mass index > 35 kg/m^2^. Written informed consent was collected from each participant.

### 2.4. Enzyme-Linked Immunosorbent Assays (ELISA) for Anti-GRP78 Autoantibodies

Recombinant human GRP78s purchased from StressMarq Biosciences or produced in the laboratory [30] were used as antigens in ELISA. The proteins were diluted to 0.5 μg/ml in PBS to coat in ELISA plates (100 μl/well) overnight at 4°C. After washing three times with PBST (PBS + 0.1% Tween 20) and blocking for 2 hours at room temperature, plasma specimens (1:100 dilution in blocking buffer) were added to the pre-coated wells to incubate for another 2 hours. After thorough washing with PBST, alkaline phosphatase-conjugated goat anti-human antibody (Invitrogen, Carlsbad, CA) was added (1:5000 dilution in blocking buffer), and color developed with p-nitrophenyl phosphate (KPL, Gaithersburg, MD). Absorbance at 405 nm was determined by a plate reader.

### 2.6. Sample Preparation for Mass Spectrometry (MS)

The various rGRP78 preparations were digested overnight with PNGase F (P7367, Sigma-Aldrich, St. Louis, MO). Following denaturation, the samples were loaded onto a 10% Bis-tris gel. The gel was stained overnight with Colloidal Coomassive. In all samples, molecular weight (MW) bands at ~78kDa were excised and digested with Trypsin Gold (Promega, Madison, WI) prior to nLCMS2 analyses as previously reported [32]. Since we observed different levels of impurities in some samples, we divided the samples into two groups based on different impurity levels (Figure 2). A sample from each of these two groups was picked at random and excised into 6 MW fractions to test the identities of the impurities.

Following trypsin digestion, the peptides were extracted, concentrated until nearly dry under vacuum, and diluted in 0.1% formic acid prior to analysis by 1-D reverse nano-scale liquid chromatographic electrospray ionization multi-stage tandem mass spectrometry (nLC-ESI-MS2).

### 2.7. Mass Spectrometry (MS)

Peptide digests from all MW fractions were analyzed by LC-nESI-MS2 in duplicate using the standard collision induced dissociation (CID), and when necessary, electron transfer dissociation (ETD) approach. While CID is very comprehensive for global proteomics studies, neutral-loss of O-linked PTMs can be helpful. Therefore, ETD mode is sometimes applied to complement CID data to maintain the weak O-linked PTMs. All runs were performed on a 1260 Infinity nano-scale high performance liquid chromatography (nHPLC) stack (Agilent, Santa Clara, CA) and separated using a 75 micron I.D. x 15 cm pulled tip C-18 column (Jupiter C-18 300 Å, 5 micron, Phenomenex, Torrance, CA). This system ran in-line with a Thermo Orbitrap Velos Pro hybrid mass spectrometer, equipped with a nano-electrospray source (Thermo Fisher Scientific, Waltham, MA). The nHPLC configuration and scan method have been described previously [32]. Of note, for the ETD experiments, samples were run on a separate nLC-ESI-MS2 system with exact same configuration as the one located at the UAB & CCC Mass Spectrometry & Proteomics Shared Facility (MSP-SF). This system was also benchmarked by the MSP-SF staff, and housed under the UAB Stem Cell Institute.

### 2.8. MS Data Conversion and Searches

The XCalibur RAW files were collected in profile mode, centroided and converted to MzXML using ReAdW v. 3.5.1. Mgf files were then created using MzXML2Search (included in TPP v. 3.5) for all scans with a precursor mass between 350Da and 2,000Da. The data was searched using SEQUEST with filters set for two maximum missed cleavages, a precursor mass window of 20ppm, trypsin digestion, variable modification C at 57.0293, and M at 15.9949. In addition, PTMs were searched as well in two separate stages for phosphorylation on Threonine (T), Serine (S), and Tyrosine (Y). Searches were performed with a curated human and *E. coli* specific subsets of the UniRef100 database, which include common contaminants such as digestion enzymes and human keratin.

### 2.9. Peptide Filtering, Grouping, and Quantification

A list of peptide IDs, generated based search results stemming from MASCOT/ MASCOT Distiller (Matrix Science, London, UK), were filtered using Scaffold Q+S (Protein Sciences, Meriden, CT) as previously described [33]. In addition, for all PTMs, we analyzed the exported Scaffold Q+S files in Scaffold PTM (Protein Sciences, Meriden, CT) where the use of assigned A-scores and localization C.I.’s allowed us to filter out potential false positive PTM assignments [34,35]. The PTMs that passed these filters were manually interpreted prior to moving forward. It is noteworthy that the enrichment step in our system, combined with 1-D PAGE, generated enough high-quality protein that further enrichment is often not necessary to obtain high confidence PTM assignments with reproducible quantification.

## 3. Results

### 3.1. Variable results in ELISA assays to measure GRP78 autoantibody levels in the study specimens

Our previous studies identified GRP78, an important regulator of the unfolded protein response [36], as an autoantigen in emphysema/COPD patients [31]. To further study the role of GRP78 autoimmunity in pathogenesis, we conducted several ELISAs to measure the concentration of plasma GRP78 autoantibodies in COPD and SC subjects. Different lots of human rGRP78 as the plate-bound antigen to detect GRP78 autoantibodies resulted in discordant ELISA findings (Figure 1A-F) while our replicate results are highly reproducible (Supplementary Figure S1). These inconsistent results led us to conduct troubleshooting assays to fully characterize potential protein contaminants, in addition to the measuring the relative purity of different lots of rGRP78. In these studies, we expressed and purified rGRP78 from *E. coli* in our laboratory, in order to minimize lot-to-lot variation of the protein antigen.

**Figure 1.**
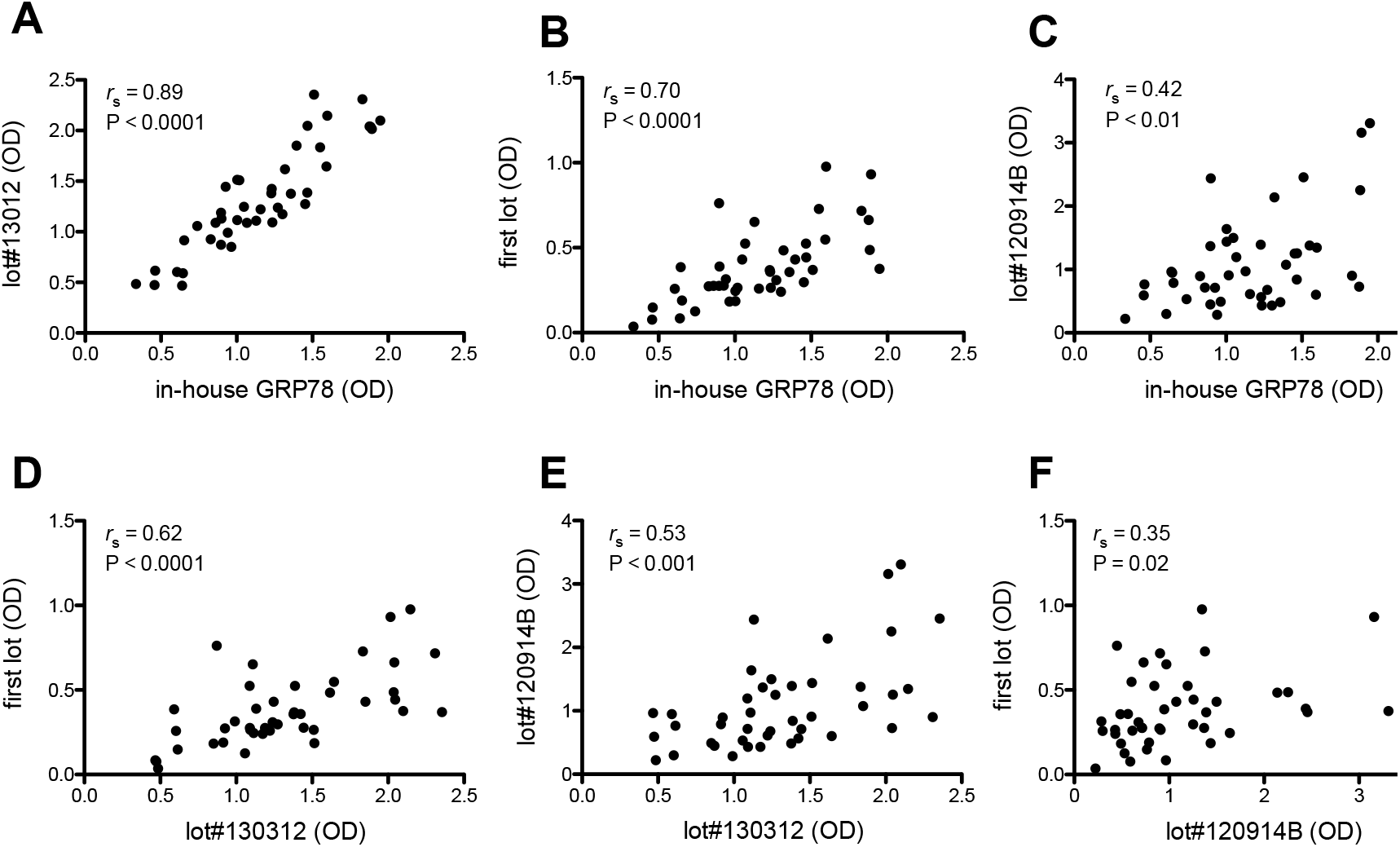
Varying ELISA results to measure GRP78 autoantibodies using different rGRP78 preparations. Dot plots show discordant associations (A-F) between levels of GRP78 IgG autoantibodies as measured by Optical Density (OD) in various ELISA assays with different lots of rGRP78 purchased commercially (first lot, lot#130312, lot#120914) and one preparation produced in our laboratory. The first lot number of commercial rGRP78 was unknown. Comparisons were made by Spearman’s rank correlation test. It should be noted that our replicate ELISA results show high level of reproducibility (Supplementary Figure S1).

### 3.2. Testing the purity and identity of different rGRP78 preparations

Different lots of commercial rGRP78 as well as the rGRP78 produced in our laboratory were run on SDS-PAGE gels and stained with Coomassie dye to separate and visualize them. Other samples were also treated with PNGase F to cleave off the N-glycans to compare the molecular weight (MW) shifts when the GRP78 proteins were de-glycosylated. SDS-PAGE gels showed most rGRP78 preparations contain the expected size (~78kDa) of human GRP78. However, different commercial lots exhibited varying levels of purity with the purest lot (lot#130312) being comparable to the rGRP78 preparation produced in our laboratory (Figure 2). In retrospect, this finding was not surprising, given the GRP78 IgG ELISA results above showed the highest degree of correlation between these two preparations (Figure 1).

**Figure 2.**
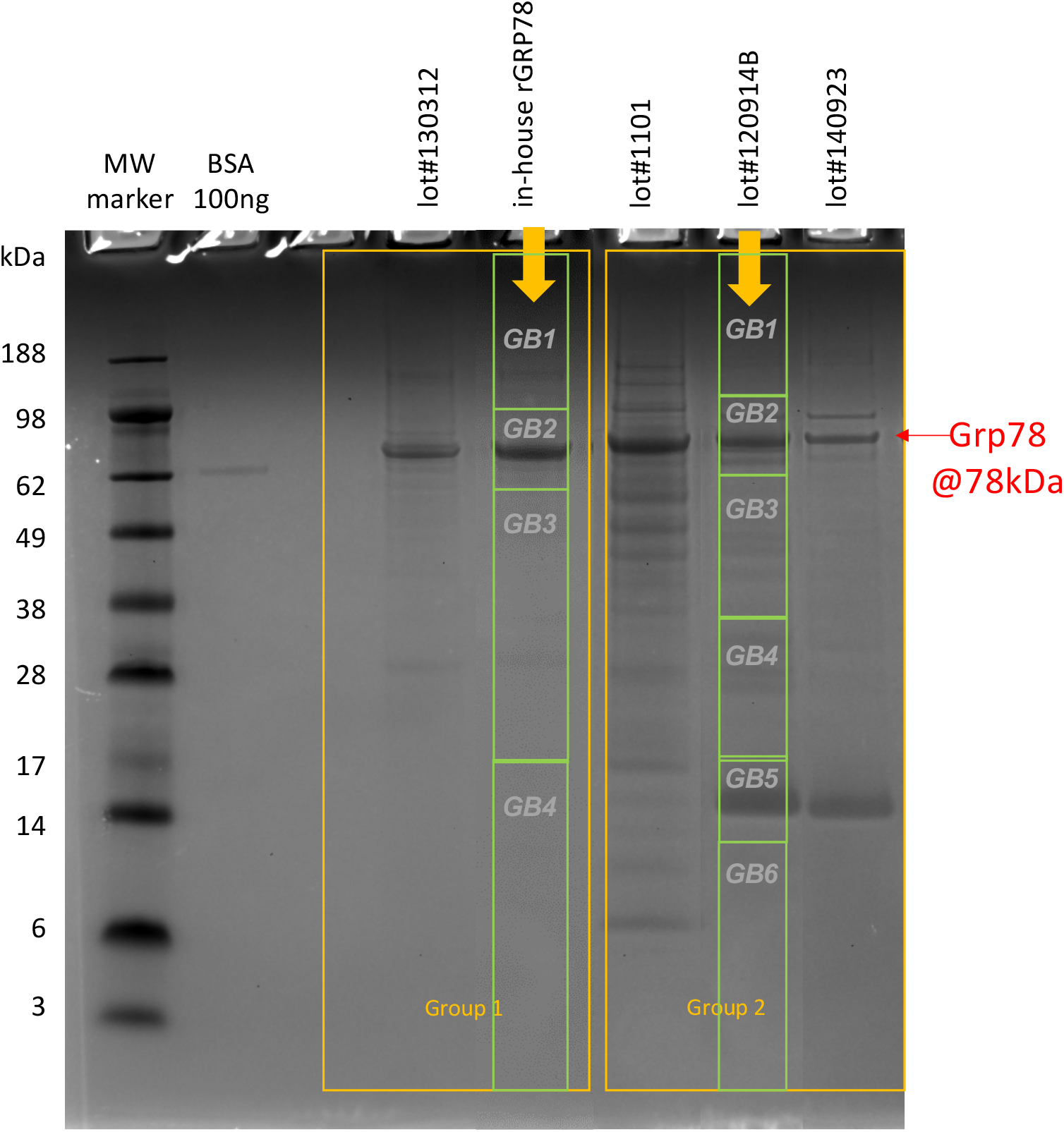
The purity of different rGRP78 samples was assessed by SDS-PAGE. Representative image of an SDS-PAGE gel showed varying levels of purity (by densitometry) among different lots of commercial rGRP78 and one sample that had been produced in our laboratory. All samples were loaded in equal amounts (5 μg) based on the results of standard BCA assays. The bands at ~78kDa were excised and digested with trypsin for MS analyses. From the densitometry results, the samples were then divided into two groups based on the level of impurities. The representative samples for each group (indicated by the arrow sign) were selected to be cut into various MW fractions (gel bands (GB) 1-6) for further LCMS2 analyses of the whole lane digestion.

MW bands at ~78kDa from all recombinant protein samples were excised and digested with trypsin prior to LCMS2 analyses. The overall GeLC-MS2 workflow for peptide/protein identification is illustrated in Figure 3. Briefly, after the MS/MS spectra were generated, we used MASCOT distiller to match each MS/MS spectra to a peptide sequence. We found that within the ~78kDa band, there were fairly high levels of purity (up to ~90% of GRP78 spectral counts in the whole lane digestion) among different rGRP78 preparations (Table 1).

**Figure 3.**
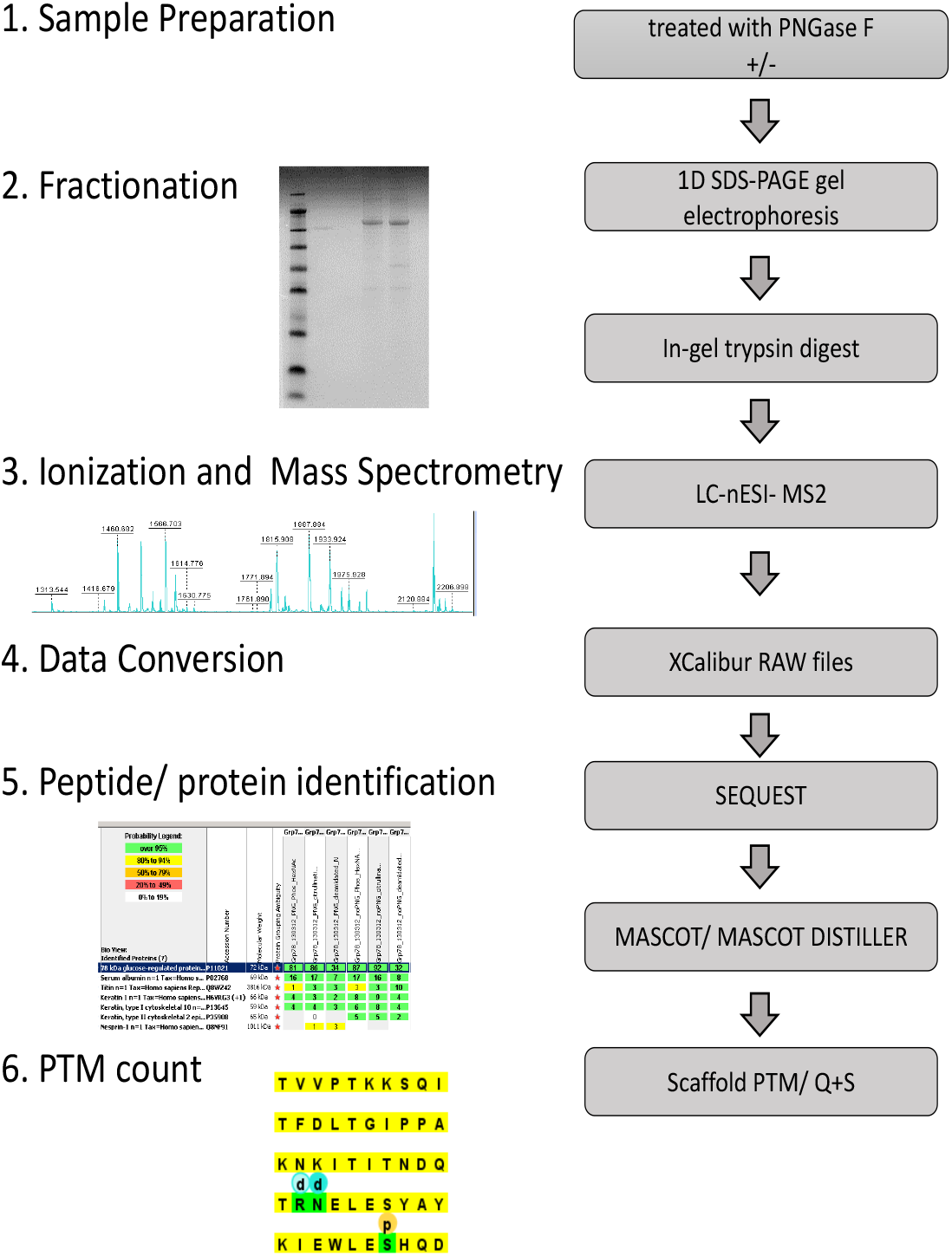
Peptide/protein and PTM identification workflow.

**Table 1.** Peptide identification within the 78kD band by spectral counting among different lots of rGRP78.

|  | lot#1101 | lot#120914B | lot#130312 | lot#140923 | in-house |
| --- | --- | --- | --- | --- | --- |
| Total Spectral Counts | 1017 | 889 | 877 | 838 | 958 |
| # GRP78 Spectral Counts | 898 | 799 | 741 | 693 | 861 |
| % GRP78/total | 88.3% | 89.9% | 84.5% | 82.7% | 89.9% |

Based on the SDS-PAGE results, we also divided the samples according to their purity levels into two groups and randomly selected one representative sample in each group to be cut into different MW fractions (as shown in Figure 2) for whole lane digestion and more comprehensive MS analyses. We found that for both groups, the various MW bands all matched to human GRP78 (Supplementary Figure S2), showing that these impurities were mostly cleaved products of the rGRP78 protein. Within a gel band, there were ~40-95% of the total spectral counts that matched to GRP78 (Supplementary Figure S2). Besides GRP78 products, minor percentages of other proteins such as Chaperonin (human), GroEL (*E. Coli),* Heat shock protein Hsp90 family (*E. Coli),* L-glutamine:D-fructose-6 phosphate aminotransferase (*E. Coli)* among others made up ~0-15% of total spectral counts in a gel band (Supplementary Figure S2).

### 3.3. Different lots of rGRP78 exhibited varying PTM levels

After peptide identification by MASCOT define, we used Scaffold PTM define/describe in Methods to determine the locations and extent of various PTMs across different preparations of rGRP78. We found slightly different quantities of phosphorylation in a number of S/T/Y sites, most notably at the T102 residue (Supplementary Figure S3 and Figure 4A). On the other hand, there were more robust changes in glycosylation amount, as inferred by changes in deamidation reaction following PNGase F treatment (Supplementary Figure S4 and Figure 4B); however, the sites/locations of these glycosylation modifications with respect to particular amino acid residue in the protein generally do not differ among different batches (Supplementary Figure S5). We also checked the native deamination levels, which were not caused by PNGase F treatment (Supplementary Figure S6) to validate that the changes in deamidations observed were specifically due to the PNGase F treatment.

**Figure 4.**
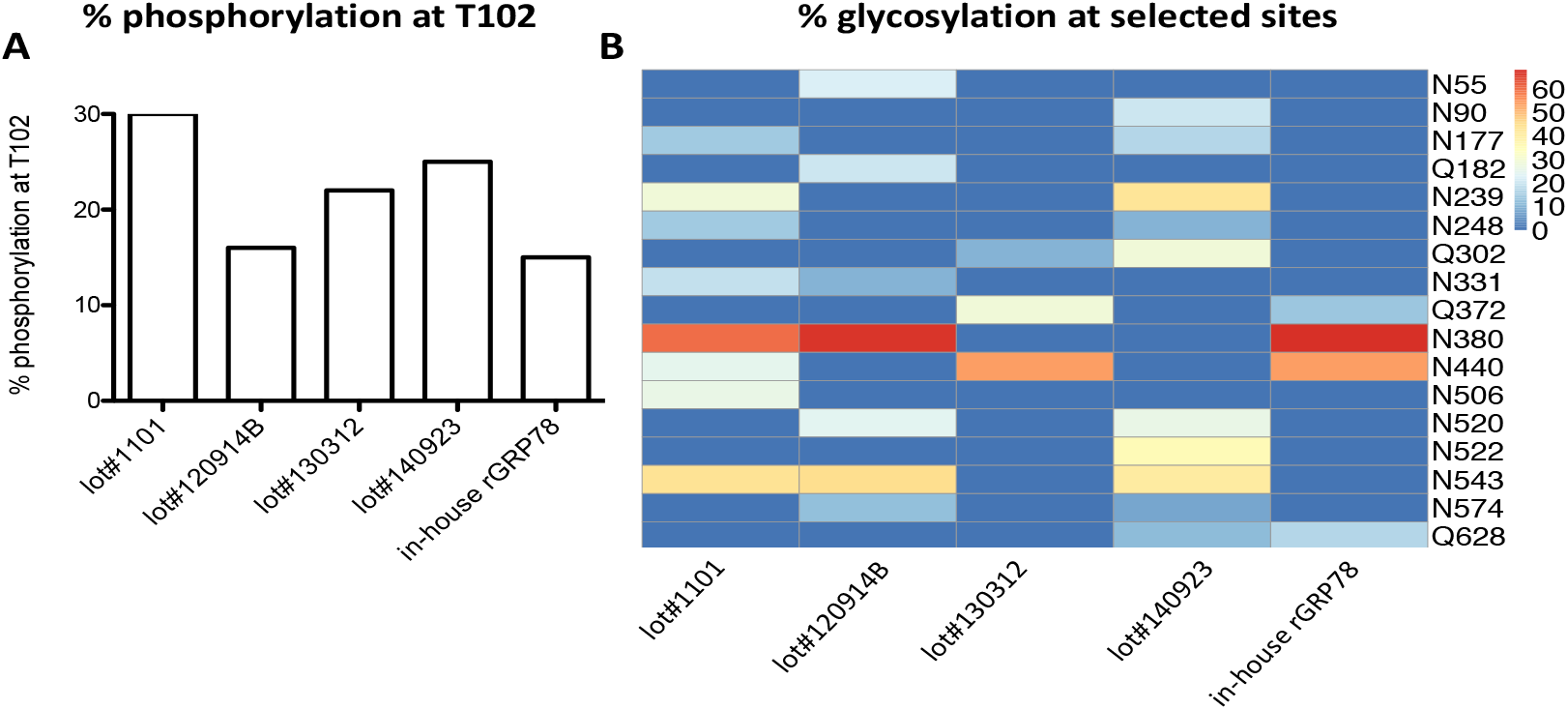
Variable PTM modifications among different lots of rGRP78. (A) The variable extent of phosphorylation on residue T102 in different preparations of rGRP78. (B) Varying levels of glycosylation modifications as inferred by changes in deamidations following PNGase F treatment.

Differences between lots were not as substantial when we searched for other PTMs such as citrullination and oxidation.

## 4. Conclusion

In troubleshooting to account for discordant ELISA results, we applied a standard GeLC-MS2 workflow, which includes the use of SDS-PAGE followed by LCMS2, to verify the quality and purity of the antigen sources, in this case: rGRP78 expressed in *E. coli.* We found varying levels of purity across each lot of rGRP78 from the same commercial product and one preparation produced in our laboratory. LCMS2 analyses showed that the majority of apparent impurities observed in the different MW bands in stained 1-D Gels were cleaved products of human GRP78. However, variation in purity levels alone cannot fully account for the discordant ELISA results, because while lot#130312 and our own lab-generated rGRP78 exhibit similar purity levels, the ELISA concordances of these two preparations are not 100%. We also found varying PTM levels in these preparations, particularly in phosphorylation and glycosylation modifications, although the locations of these PTMs with respect to the particular amino acid residues in the protein do not differ. Therefore, besides varying purity levels, changes in PTMs potentially lead to inconsistent results observed in our immunoassays.

In this report, we also presented a standard workflow to assist as a guideline in the testing for the purity and PTM levels of recombinant protein products to help researchers perform QC analyses, and when applicable, troubleshoot problematic laboratory immunoassays. Our report is intended to raise awareness of variations within batches of “purified” proteins and suggests that such samples are routinely tested to ensure the reproducibility and rigor of scientific research applications.

## Supplementary Materials

Figure S1: High level of reproducibility in a representative ELISA assay to measure GRP78 autoantibodies in COPD plasma; Figure S2: Protein ID’s with % of total spectral counts/gel band; Figure S3: The fractions of GRP78 peptides with phosphorylation modification from various rGRP78 preparations; Figure S4A-E: The fractions of GRP78 peptides with glycosylation modifications (as inferred from the changes in deamidation levels) from various rGRP78 preparations; Figure S5: An example of a side-by-side comparison between the locations of various PTMs; Figure S6A-B: The fractions of GRP78 peptides with native deamidation modification (i.e. without PNGase F treatment) in different rGRP78 preparations.

