## Supplementary Figure S1 for "A Method for Checking Recombinant Protein Quality to Troubleshoot for Discordant Immunoassays"

### Slide 1
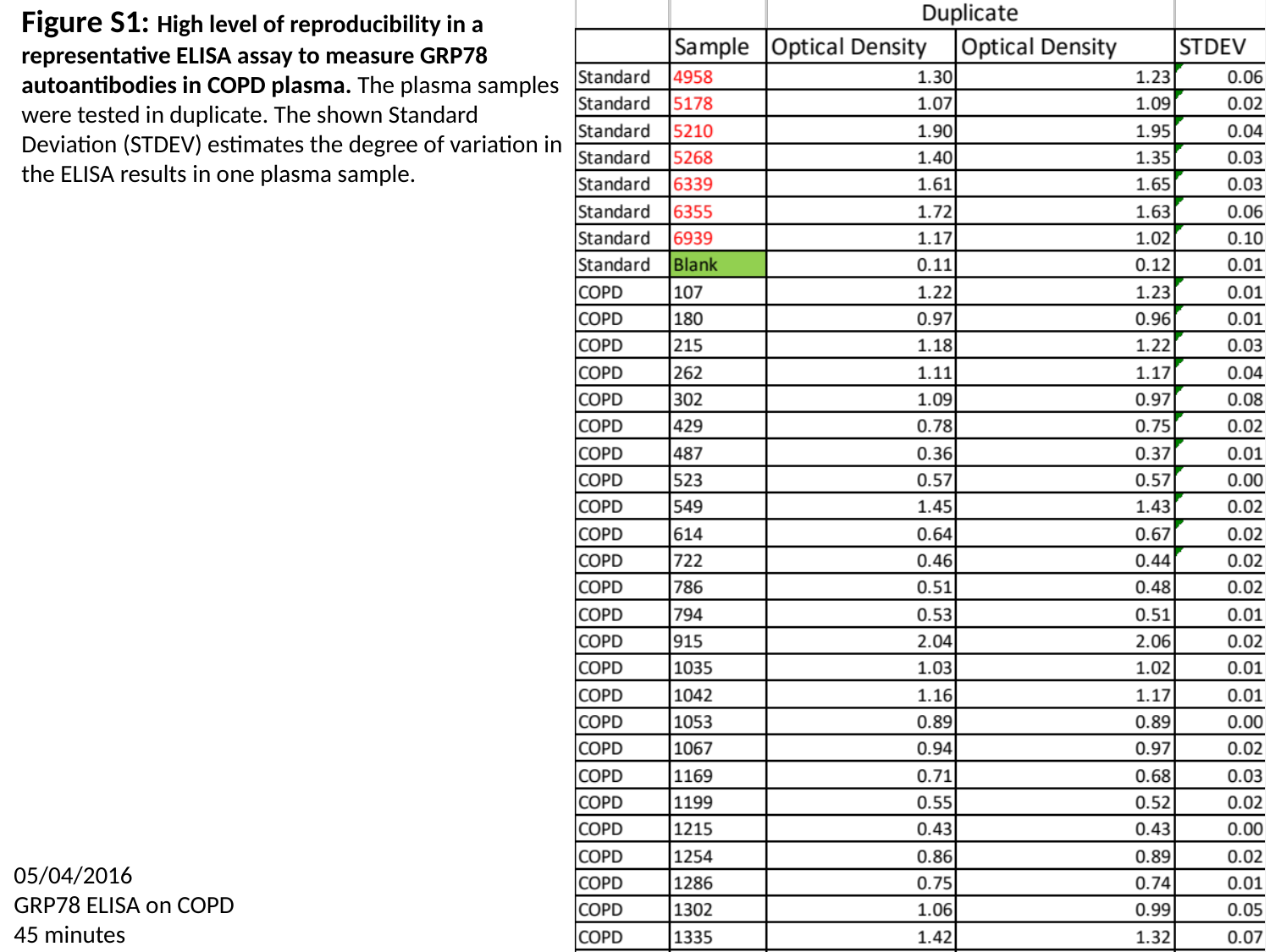

Figure S1: High level of reproducibility in a
representative ELISA assay to measure GRP78
autoantibodies in COPD plasma. The plasma samples
were tested in duplicate. The shown Standard
Deviation (STDEV) estimates the degree of variation in
the ELISA results in one plasma sample.
05/04/2016
GRP78 ELISA on COPD
45 minutes
