## Supplementary Figure S2 for "A Method for Checking Recombinant Protein Quality to Troubleshoot for Discordant Immunoassays"

### Slide 1
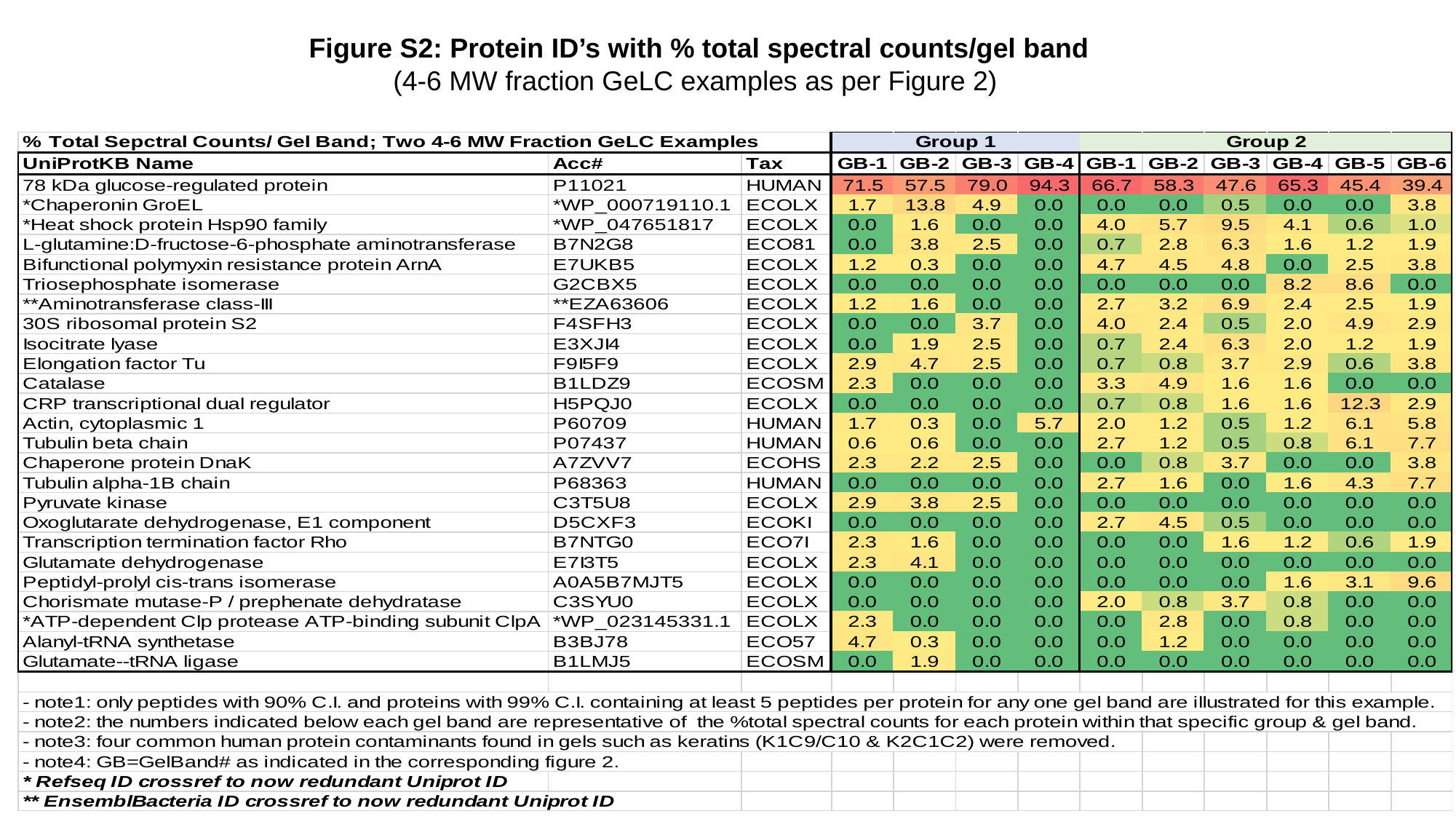

Figure S2: Protein ID’s with % total spectral counts/gel band
(4-6 MW fraction GeLC examples as per Figure 2)
