## Supplementary Figure S4 for "A Method for Checking Recombinant Protein Quality to Troubleshoot for Discordant Immunoassays"

### Slide 1
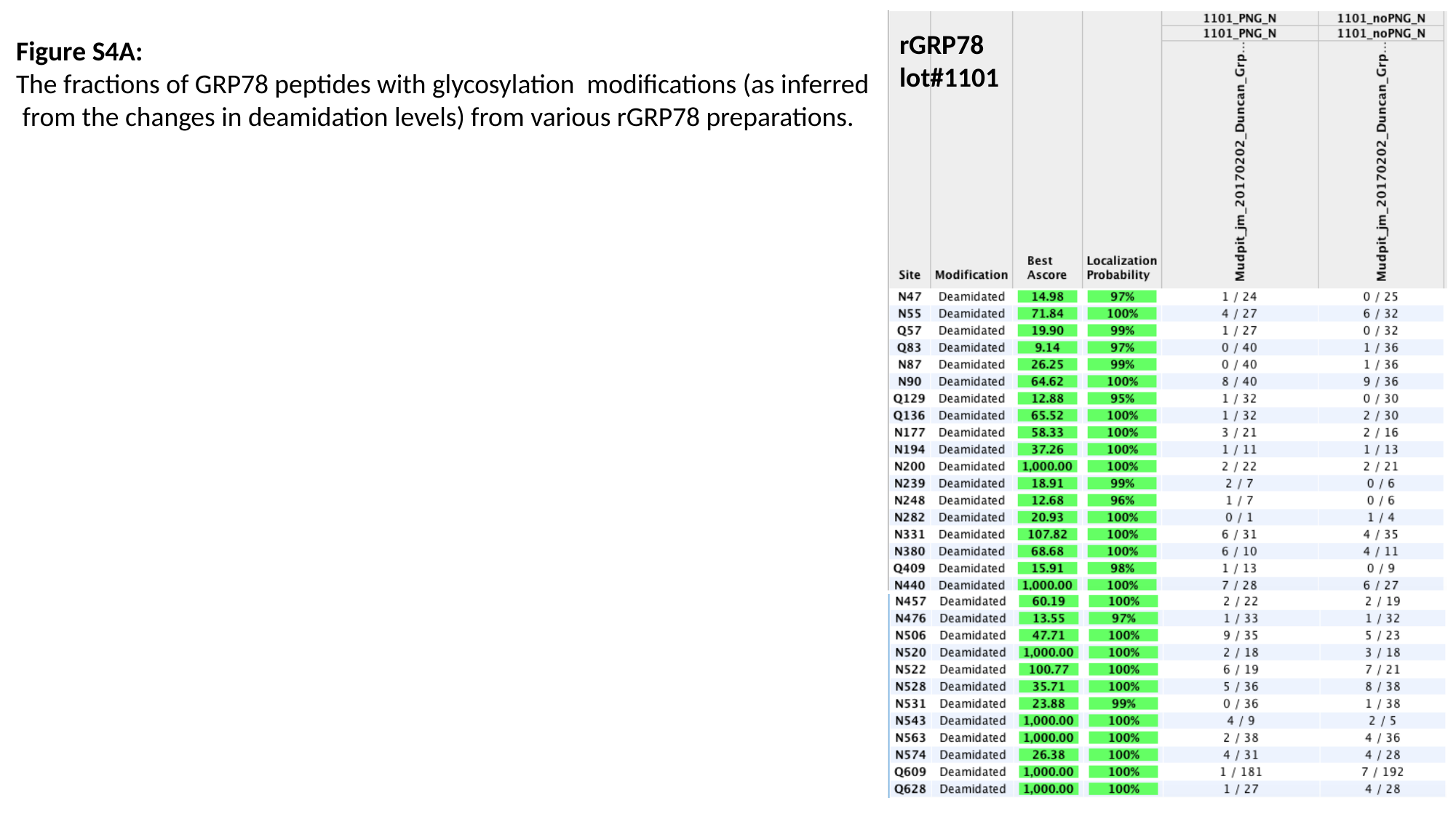

Figure S4A:
The fractions of GRP78 peptides with glycosylation modifications (as inferred
 from the changes in deamidation levels) from various rGRP78 preparations.
rGRP78
lot#1101

### Slide 2
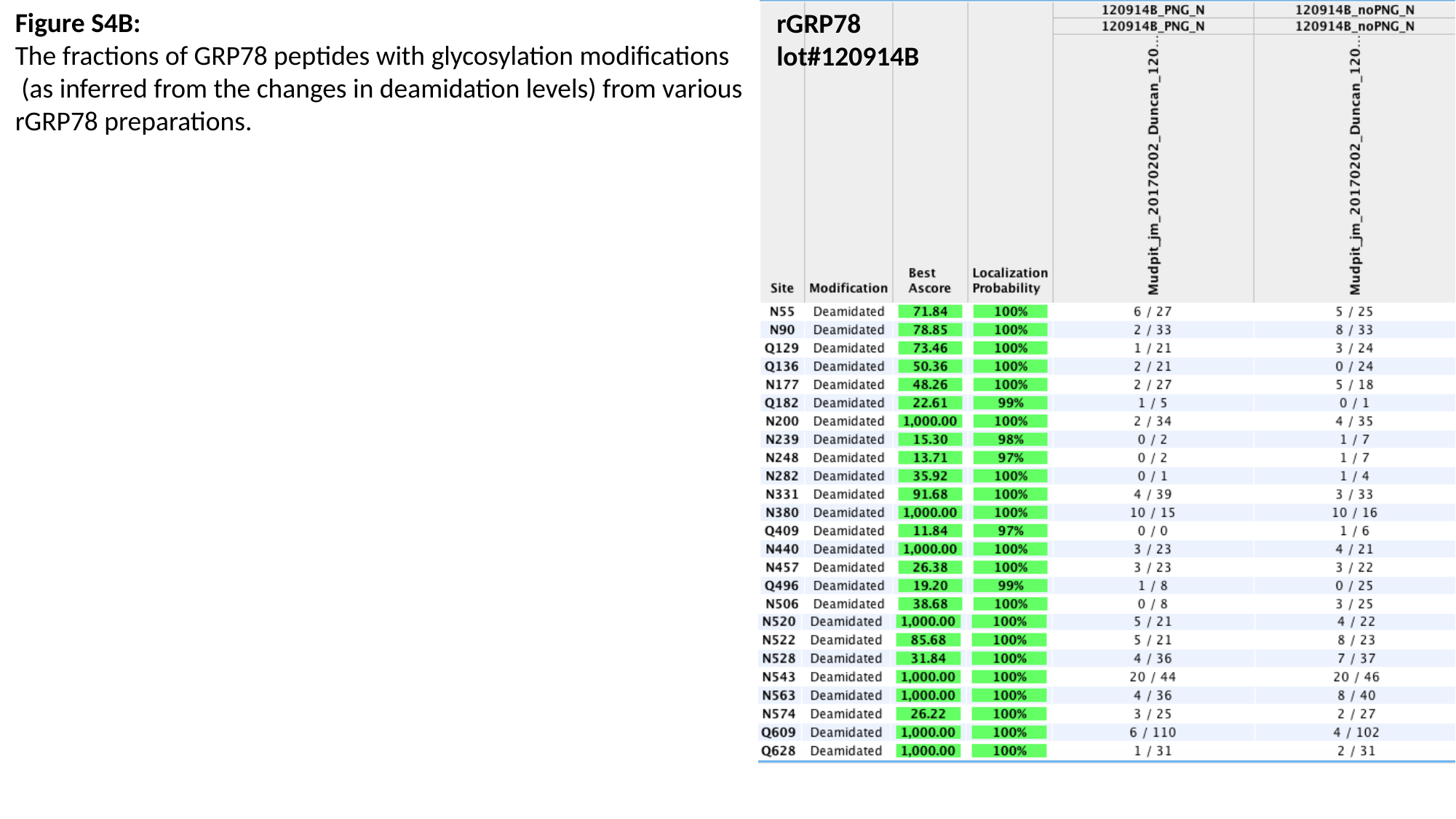

Figure S4B:
The fractions of GRP78 peptides with glycosylation modifications
 (as inferred from the changes in deamidation levels) from various
rGRP78 preparations.
rGRP78
lot#120914B

### Slide 3
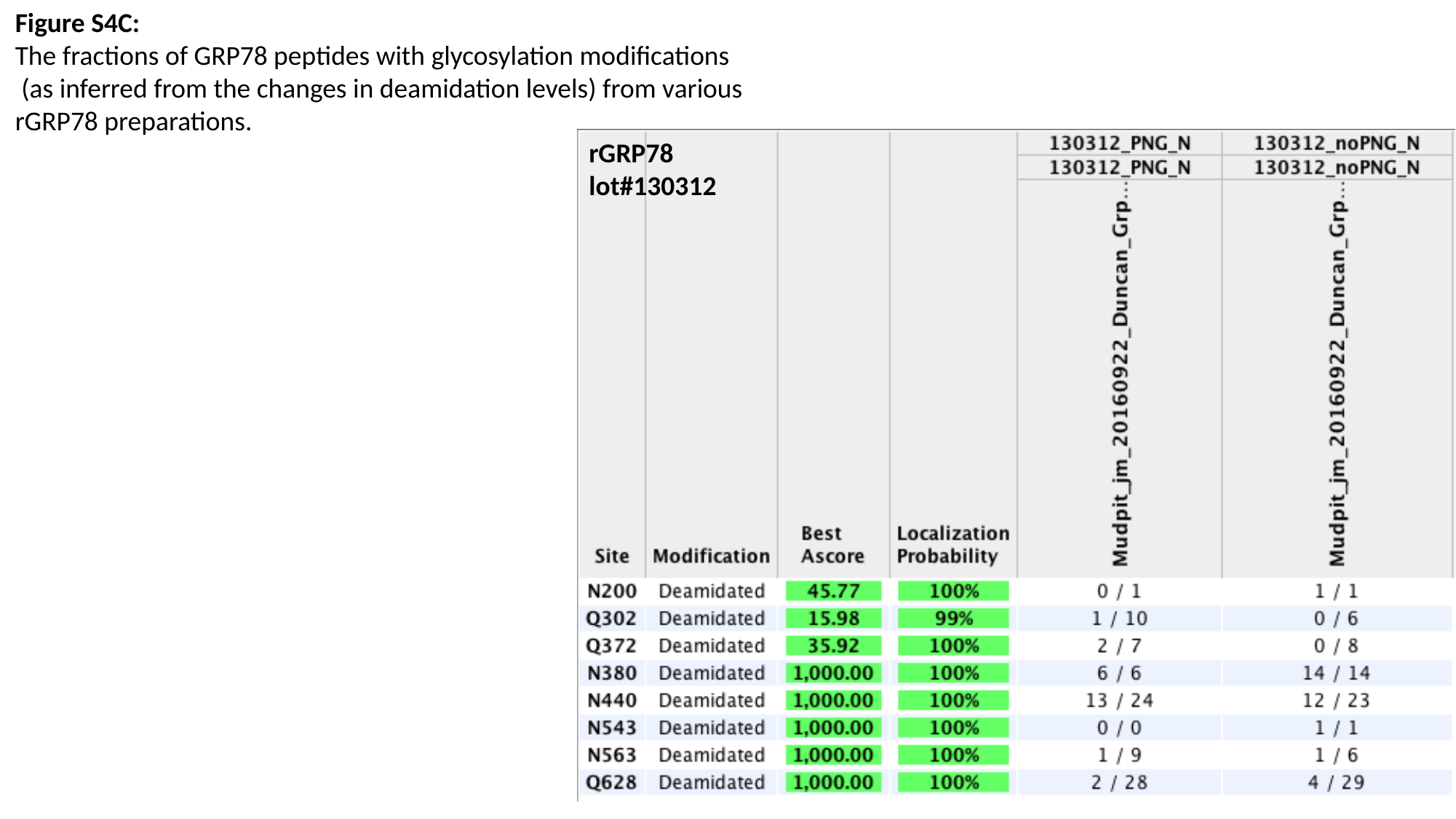

Figure S4C:
The fractions of GRP78 peptides with glycosylation modifications
 (as inferred from the changes in deamidation levels) from various
rGRP78 preparations.
rGRP78
lot#130312

### Slide 4
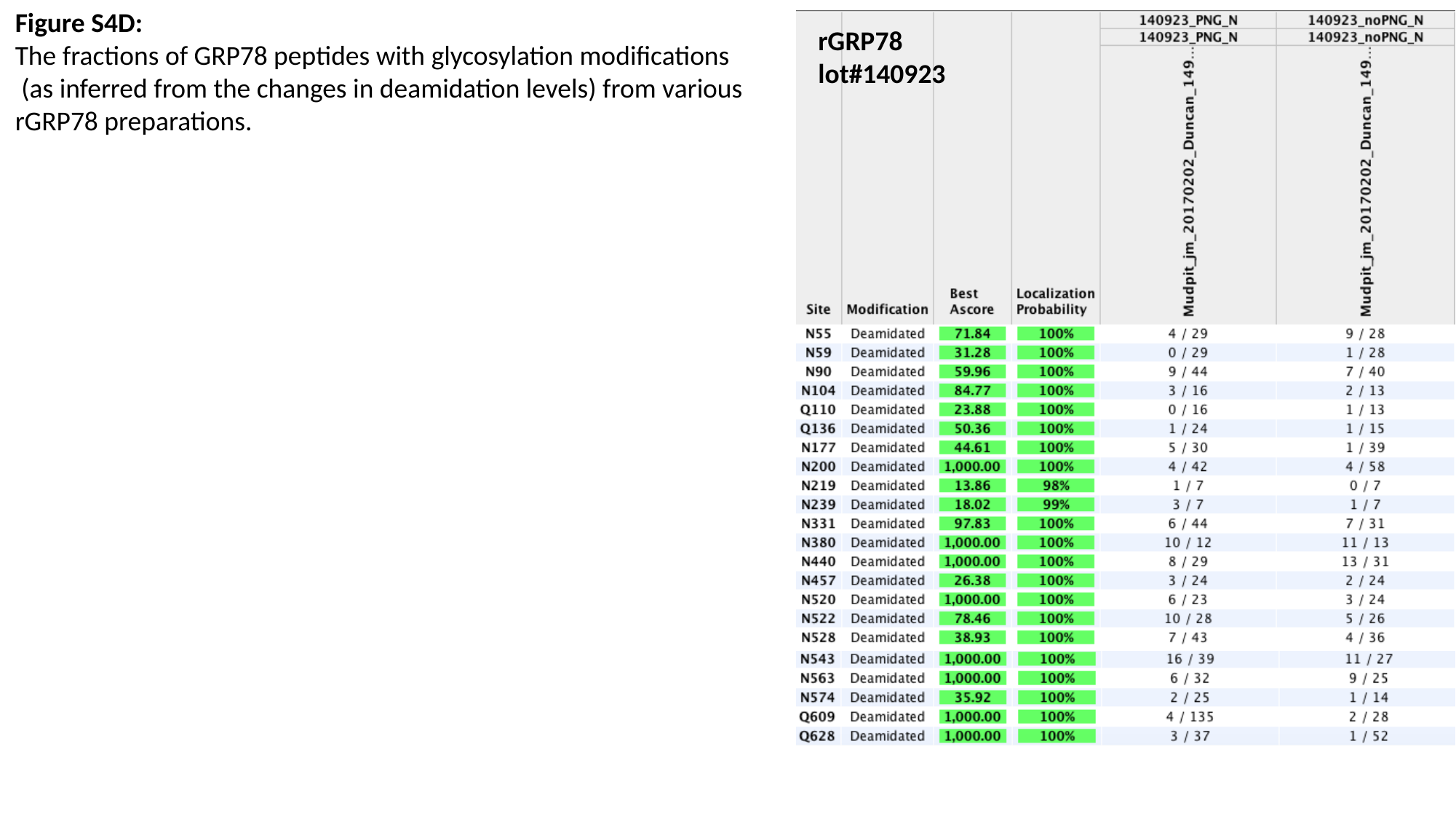

Figure S4D:
The fractions of GRP78 peptides with glycosylation modifications
 (as inferred from the changes in deamidation levels) from various
rGRP78 preparations.
rGRP78
lot#140923

### Slide 5
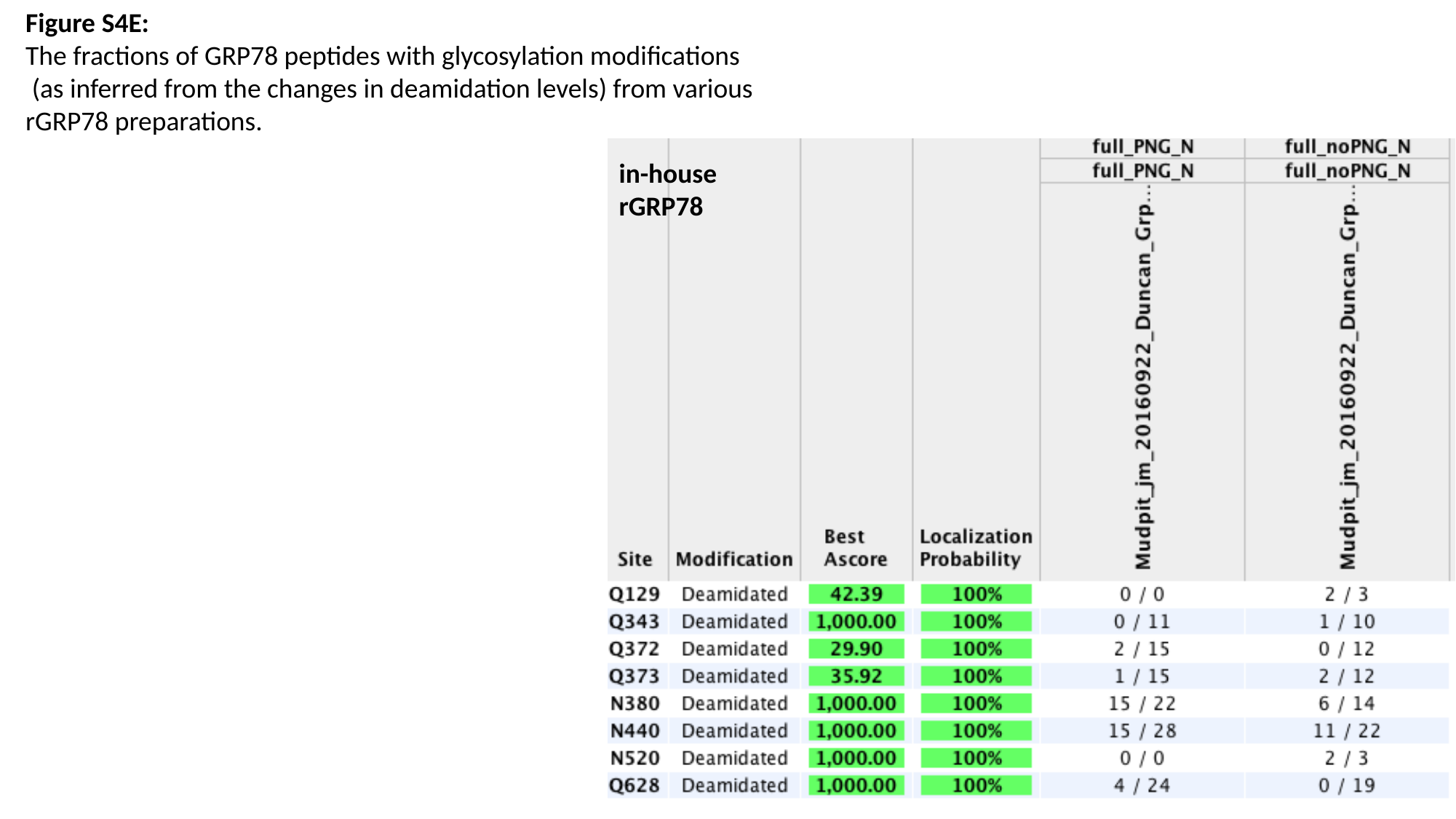

Figure S4E:
The fractions of GRP78 peptides with glycosylation modifications
 (as inferred from the changes in deamidation levels) from various
rGRP78 preparations.
in-house
rGRP78
