## Supplementary Figure S3 for "A Method for Checking Recombinant Protein Quality to Troubleshoot for Discordant Immunoassays"

### Slide 1
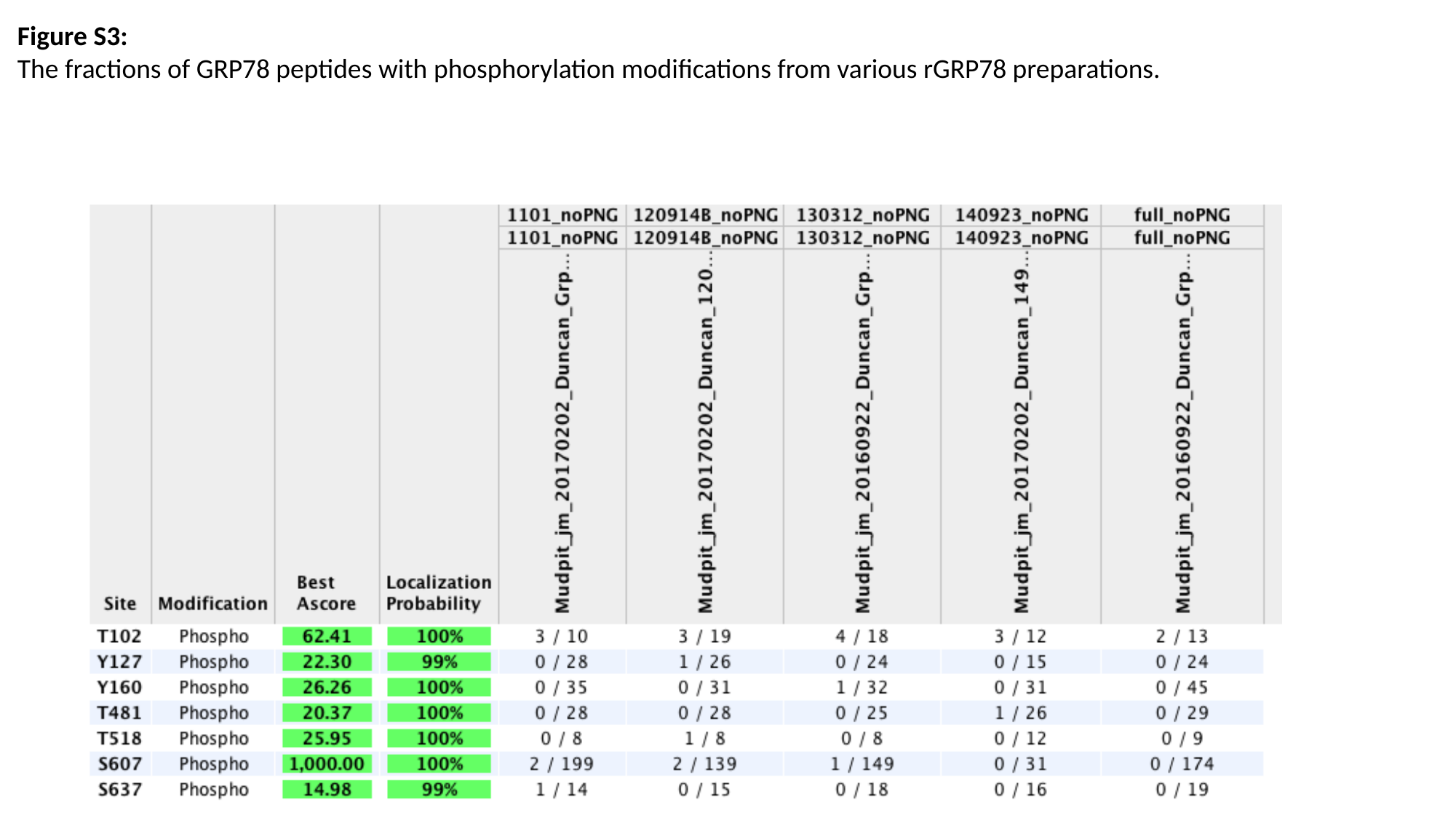

Figure S3:
The fractions of GRP78 peptides with phosphorylation modifications from various rGRP78 preparations.
