## Supplementary Figure S5 for "A Method for Checking Recombinant Protein Quality to Troubleshoot for Discordant Immunoassays"

### Slide 1
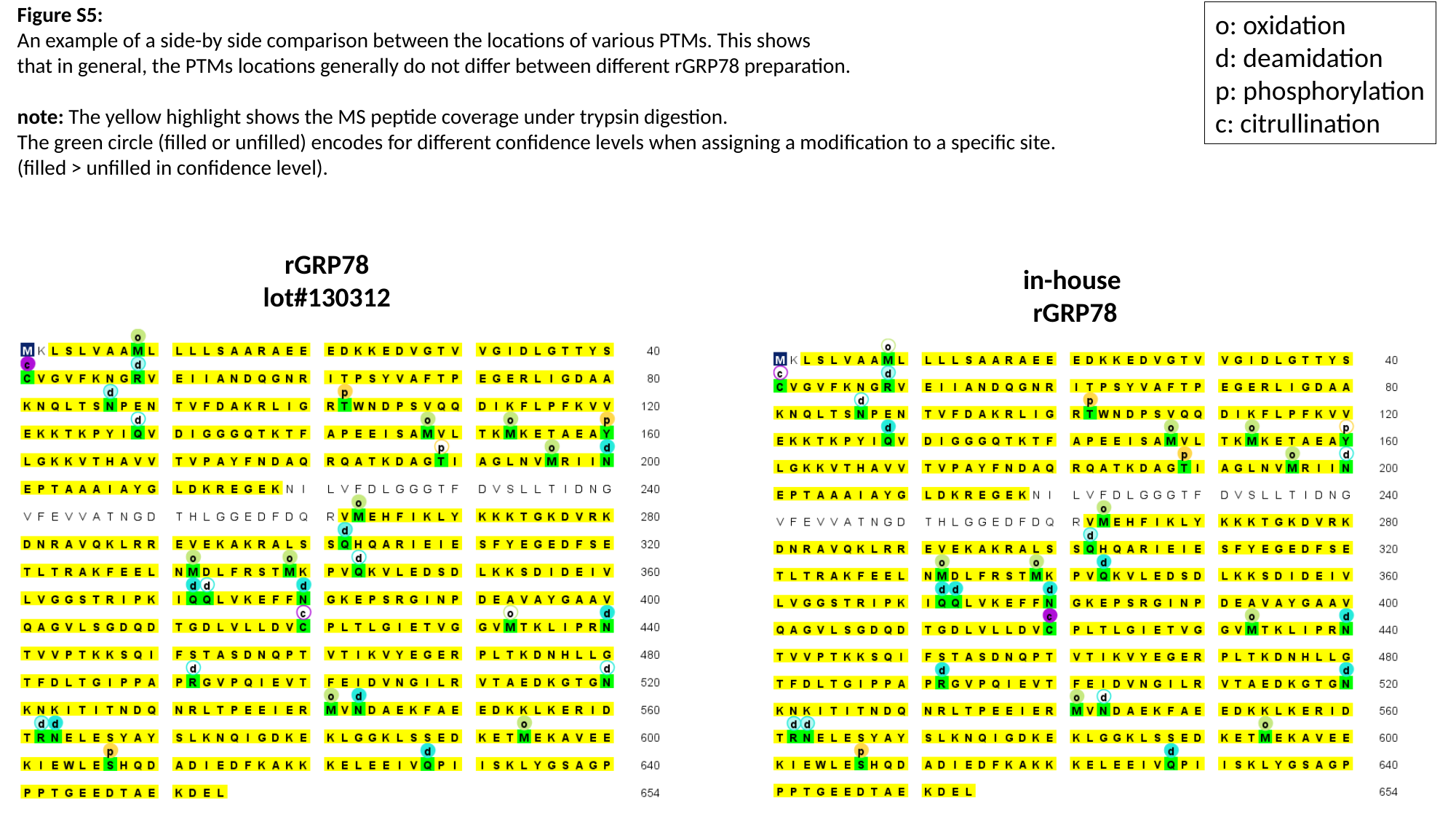

Figure S5:
An example of a side-by side comparison between the locations of various PTMs. This shows
that in general, the PTMs locations generally do not differ between different rGRP78 preparation.
note: The yellow highlight shows the MS peptide coverage under trypsin digestion.
The green circle (filled or unfilled) encodes for different confidence levels when assigning a modification to a specific site.
(filled > unfilled in confidence level).
o: oxidation
d: deamidation
p: phosphorylation
c: citrullination
rGRP78
lot#130312
in-house
rGRP78
