## Supplementary Figure S6 for "A Method for Checking Recombinant Protein Quality to Troubleshoot for Discordant Immunoassays"

### Slide 1
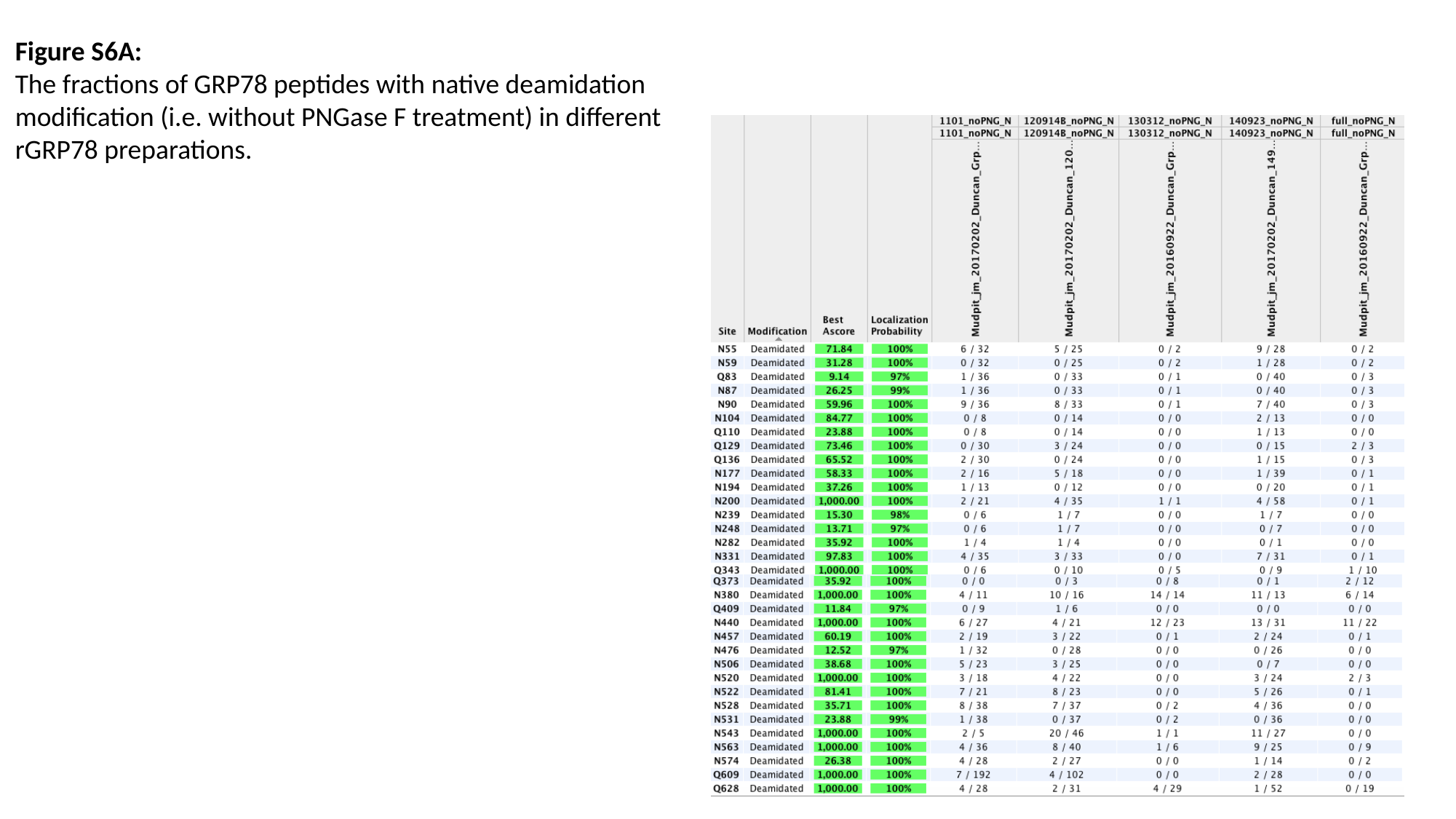

Figure S6A:
The fractions of GRP78 peptides with native deamidation
modification (i.e. without PNGase F treatment) in different
rGRP78 preparations.

### Slide 2
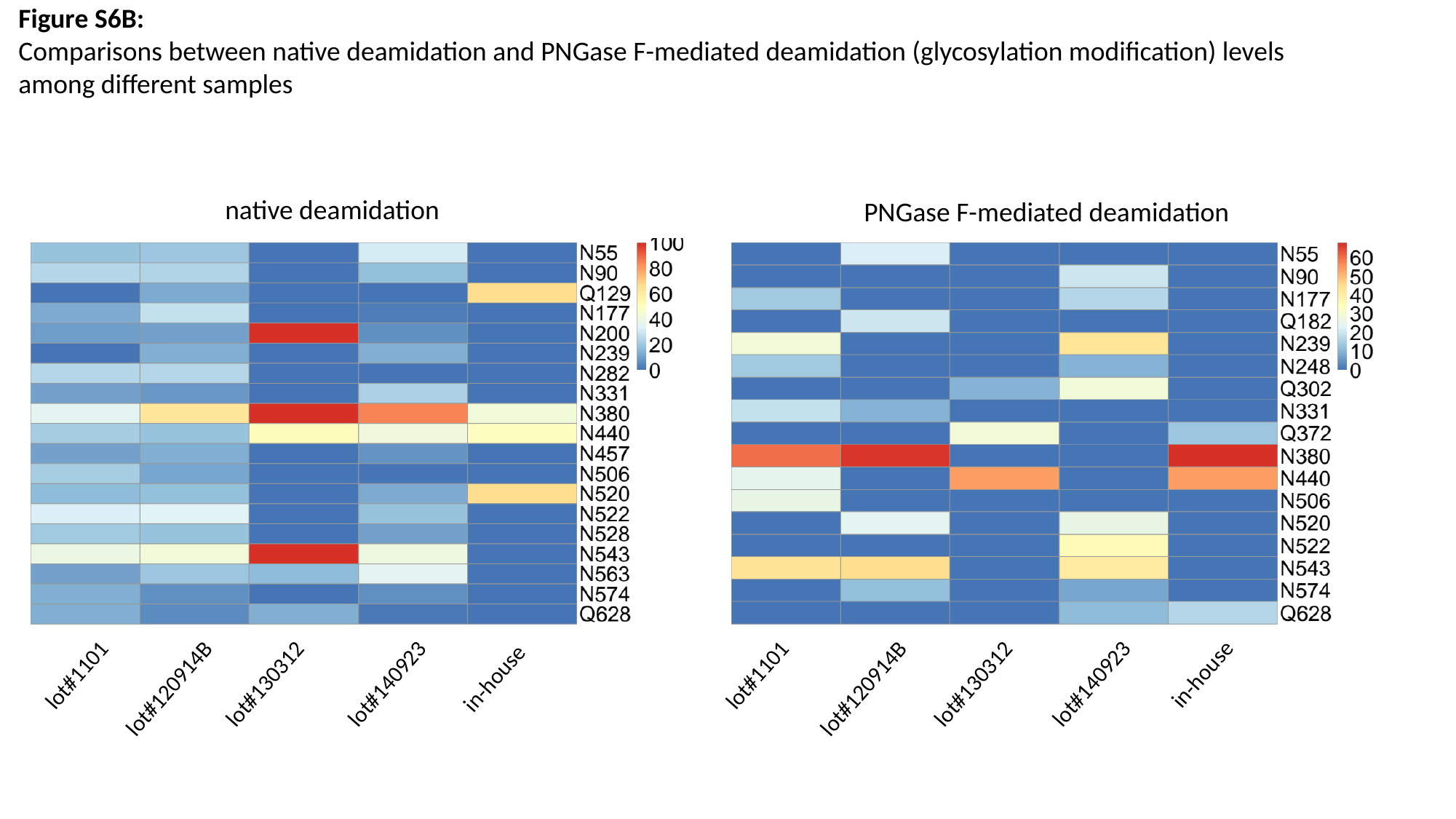

Figure S6B:
Comparisons between native deamidation and PNGase F-mediated deamidation (glycosylation modification) levels
among different samples
native deamidation
PNGase F-mediated deamidation
in-house
lot#1101
lot#1101
in-house
lot#130312
lot#140923
lot#130312
lot#140923
lot#120914B
lot#120914B
